# ganon2: up-to-date and scalable metagenomics analysis

**DOI:** 10.1101/2023.12.07.570547

**Authors:** Vitor C. Piro, Knut Reinert

## Abstract

The fast growth of public genomic sequence repositories greatly contributes to the success of metagenomics. However, they are growing at a faster pace than the computational resources to use them. This challenges current methods, which struggle to take full advantage of massive and fast data generation. We propose a generational leap in performance and usability with ganon2, a sequence classification method that performs taxonomic binning and profiling for metagenomic analysis. It indexes large datasets with a small memory footprint, maintaining fast, sensitive, and precise classification results. Based on the full NCBI RefSeq and its subsets, ganon2 indices are on average 50% smaller than state-of-the-art methods. Using 16 simulated samples from various studies, including the CAMI 1+2 challenge, ganon2 achieved up to 0.15 higher median F1-Score in taxonomic binning. In profiling, improvements in the F1-Score median are up to 0.35, keeping a balanced L1-norm error in the abundance estimation. ganon2 is one of the fastest tools evaluated and enables the use of larger, more diverse, and up-to-date reference sets in daily microbiome analysis, improving the resolution of results. The code is open-source and available with documentation at https://github.com/pirovc/ganon.

## 1 Introduction

The study of the collective complete genomic content from environments bypassing the cultivation of clonal cultures is known as metagenomics. Through DNA extraction and sequencing, the collection of microorganisms in an environment can be profiled in terms of taxonomic content, abundance, and, consequently, their function [1]. Metagenomics enables the analysis of virtually any community, ranging from sewage, subway systems, soil, oceans, and the human gut, to cite a few. Applications are limitless: clinical, industrial, and agricultural; effects of medication, diet, and lifestyle; antimicrobial resistance; pathogen detection; outbreak investigation; bioforensics; etc.

Computational metagenomics enables these applications through automated methods and data analysis [2]. However, analyzing communities is not trivial due to their complexity, high diversity, and limitations of the available technologies to translate genomic content into understandable and error-free data. Therefore, computational metagenomics faces several methodological challenges. One of the most critical aspects of current analysis is the data complexity and size of the data.

First, obtaining sequences from unknown environments through high-throughput sequencing equipment requires deep sequencing to detect low-abundant organisms, thus producing massive data sets. Large-scale studies can generate hundreds or even thousands of samples in high throughput (millions to billions of base pairs sequenced per sample), amplifying the issue. Second, the size of assembled genomic reference sequence repositories is growing exponentially [3]. However, they are invariably necessary in the analysis process of most metagenomics studies. This growth is closely related to the decreased cost of sequencing that has outpaced Moore’s law [4] since 2008. This means that data are being generated at a much faster rate than the availability of computational resources to analyze it. Specialized tools and algorithms are therefore essential to scale with massive datasets, which usually require high-performance computational resources. The goal is to enable metagenomic analyzes to scale to larger sets of data, using a limited amount of memory and in manageable time (minutes to hours per sample) while keeping sensitivity and precision high.

Due to the decrease in sequencing costs and improvements in related technologies and algorithms, large amounts of assembled genomic data are being generated and constantly deposited in public data repositories [5]. With large-scale studies, some very well-studied communities are now mostly covered by reference sequences of well-characterized species (e.g. the human gut microbiome). However, reference data is not equally distributed in the tree of life [6]. Well-described biomes represent a very small fraction of the still unknown microbial world living and evolving around us [7]. Extreme data growth is mainly biased by human-related bacteria and model organisms (e.g. *Escherichia coli*).

To extract the most information from new clinical or environmental data, it is essential to be up-to-date with this exponential growth of assembled genome sequences publicly available [8]. NCBI RefSeq [6], a common resource for metagenomics reference data, had in the last 15 years (November 2009 to November 2024) an annual growth rate of around 20% in the number of species. However, there is an imbalanced representation in genomic repositories. For example, in NCBI RefSeq, the 187 most represented species in the Archaeal, Bacterial, Fungal, and Viral Complete Genomes subsection have as many base pairs as the remaining 27662 species (as of April 2024). In the full RefSeq, the ratio is 82 to 87979. Moreover, the lack of metadata and clear documentation makes data selection a difficult and error-prone step. This leads most studies to use the default, outdated, or pre-built databases, leaving behind a large amount of information. For example, a subset of RefSeq Complete Genomes has approximately only 37% of the diversity in terms of number of species compared to the full RefSeq (Table 1). Another example: a 2-year-old outdated full RefSeq release (November 2022) has 34208 fewer species than the actual release (November 2024).

**Table 1.** Reference sets used in this article. Data obtained in 2024-04-20 for archaea, bacteria, fungi, and viral groups. RefSeq CG+RG is the union of the complete and reference genome subsets. CG and RG sets have common references, so the values are not an exact sum of both sets.

|  | # sequences | # assemblies | # species | Mbp |
| --- | --- | --- | --- | --- |
| **RefSeq** | 36725285 | 366941 | 64616 | 18767.31 |
| **RefSeq CG** | 110470 | 55114 | 24238 | 2173.06 |
| **RefSeq RG** | 1083138 | 19890 | 19888 | 1306.45 |
| **RefSeq CG+RG** | 1183909 | 69600 | 37864 | 3195.85 |

This information is difficult to track, and it is up to the researcher to mine those repositories. Extensive studies on the subject confirmed the strong improvement in metagenomic classification with larger reference sequence databases [8, 9]. Other studies confirm the importance of reference choice for research outcomes [10–12]. The use of an outdated version of references or the inability to use all currently available references leads most methods, in the best case, to underperform and, mostly common, to generate inconsistent, wrong, or incomplete results.

To address the aforementioned challenges, we propose ganon2, a sequence classification and metagenomics profiling tool that is able to classify large amounts of sequences against large amounts of references in a short time, using low amounts of memory with high sensitivity and precision. ganon2 builds on its first version, and it provides a generational leap in terms of computational performance and results. ganon2 is open source and available at: https://github.com/pirovc/ganon

## 2 Methods

### 2.1 ganon2

ganon2 improves and builds on its first version [13] as an analytical tool for metagenomics analysis. It ships an efficient algorithm for sequence classification with the Hierarchical Interleaved Bloom Filter (HIBF) [14, 15] from the SeqAn3 [16] library with several pre- and post-processing procedures to facilitate and improve its use in metagenomics analysis. ganon2 includes an integrated step for better database selection, download, and construction using standard (RefSeq or GenBank) or custom reference sequences, pairing them with NCBI or GTDB taxonomy, with support for assembly, strain, or sequence-level analysis. It further processes classification results to provide either sequence or taxonomic profiles, with abundance estimation including correction for genome sizes, multimatching read reassignment with the Expectation-Maximization (EM) and/or the Lowest Common Ancestor (LCA) algorithm with multiple reporting filters and outputs for better integration into downstream analyzes. ganon2 is a user-friendly software that automates most of its processes and provides a data-based optimized set of parameters and clear trade-offs. Thus, it helps researchers focus on their results and analysis rather than on data handling and tool execution.

#### 2.1.1 Indexing and database build

ganon2 uses the HIBF [15] as its main index, a set membership data structure composed of several interconnected Interleaved Bloom Filters (IBF) [17]. IBFs are a collection of interleaved bit-by-bit bloom filters [18].

HIBF is a fast and space efficient data structure that is designed to perform well with unbalanced data sets, which are commonly used in metagenomics analysis. As an example, the number of publicly available genome sequences in NCBI RefSeq is very unbalanced, with better-represented human pathogens and well-studied reference organisms [6].

The IBF is built on the concept of user bins, which are sets of interest to be indexed and queried. For example, a user bin can be a set of genome sequences of a certain species group. An index is formed by several user bins that have to be of the same size to be interleaved [17]. The bigger the bins, the more space an index requires. The more bins, the longer it will take to query it. Therefore, a balance between the number of bins and their size is crucial. The HIBF builds on this concept and extends the IBF usage by splitting large user bins and merging small user bins into internal technical bins to better accommodate their contents and achieve optimal use of space. User bins that were merged into one technical bin require an additional and smaller IBF structure to define their membership. The smaller IBFs are linked in a hierarchical data structure, forming an HIBF ([15] Figure 2).

ganon2 creates indices based on a set of representative k-mers for each user bin. They are optionally transformed using winnowing minimizers to reduce space requirements, at a cost of sensitivity in classification. If used in index construction, the same transformation is applied when querying. ganon2 supports the IBF and HIBF indices and builds them based on a maximum false positive rate value, k-mer size, window size, and number of hash functions for the bloom filter. User bins can be set to many possible targets: taxonomic ranks (e.g., genus, species, etc.), strain/assembly level, file or sequence level, and custom specialization. ganon2 directly links the contents of the bins to a taxonomy of choice (NCBI, GTDB, or custom) which are later used to annotate and post-process the raw query results. Additionally, approximate genome sizes are calculated for each taxon, which is later used to normalize matches and estimate relative abundances.

#### 2.1.2 Classification

ganon2 follows the assumption that the more k-mers shared between a read and a reference, the better the match. To control matches between reads and references, two thresholds are used: cutoff and filter. The cutoff threshold controls the strictness of the matching algorithm, setting the minimum percentage of k-mers (or minimizers) of a read to be shared with a reference to consider a match. The filter threshold is relative to the best and worst scoring match after the cutoff and controls the percentage of additional matches (if any) that will be reported, sorted from the best to worst. The filter threshold will not change the total number of matched reads, but controls the amount of unique or multi-matched reads. An additional filter is applied at the end to control for the false positive rate of the query [19], removing possible spurious matches based on the false positive rate of the database.

After classification, ganon2 post-processes the result to solve multiple matching reads using the lowest common ancestor (LCA) [20] and/or expectation maximization (EM) algorithm [21]. The EM algorithm will re-assign reads with multiple matches among the possible targets based on the distribution of uniquely matching reads. This is done iteratively until the re-assignments converge and do not change. Alternatively, the LCA algorithm will re-assign reads with multiple matches to lowest common ancestors in the taxonomic tree.

### 2.2 Evaluations

ganon2 was evaluated in terms of sequence classification (binning) and abundance estimation (profiling), both using taxonomy for evaluation. Although similar, those two evaluations should be independently assessed. In binning, the ability to assign each read or sequence to its correct origin is evaluated. In profiling, the ability to detect members of a sample and their respective abundance is evaluated [22]. Binning and profiling are also evaluated with different metrics and supported by different tools.

We first compare ganon2 (v2.1.0) with similar methods evaluating not only their results but also the speed and memory consumption for database creation and classification. We evaluated: kmcp (v0.9.4) [23], kraken2 (v2.1.3) [24], bracken (v2.9) [25], and metacache (v2.4.2) [26]. Metacache was evaluated only for binning, as it does not perform taxonomic profiling. Those methods were selected based on the following criteria: open-source code, actively maintained and/or highly used and cited, scalable to handle very large data in terms of database construction and sequence classification, execution in doable time (hours/few days for building, minutes/hours per sample), possibility to create custom databases with nucleotide sequences, ability to perform taxonomic binning and/or profiling.

To evaluate ganon2 and the aforementioned methods, we use the RefSeq database in 4 different configurations: (1) full, (2) complete genomes (CG), (3) reference genomes (RG), and (4) complete and reference genomes (CG+RG) (Table 1). Note that reference genome terminology has been used since the end of 2024 for previously called representative genomes [27]. RefSeq is a commonly used source for performing metagenomics analysis and provides a good standard for reference sequences of high quality and broad diversity. RefSeq RG contains just one selected assembly for each species but with a higher number of sequences in comparison to the RefSeq CG, some of them of lower quality with fragmented contigs. RefSeq CG is the most commonly used subset, represented only by genomes marked as complete, providing a theoretically higher quality. The full RefSeq has the highest diversity (64616 species) and includes both CG and RG, being almost 9 times larger in raw size than the RefSeq CG. We further evaluated similar subsets with the GenBank database and, although it contains higher diversity, it achieved the overall worst results in our evaluations. ganon2 automates the download, build, and update of databases based on RefSeq and GenBank and any of their possible subsets (reference or complete genomes, by organism group or taxa, top assemblies). RefSeq provides a very generalized set of possible genomes to analyze metagenomics data. However, in studies from well-characterized environments (e.g. human gut), a customized database is recommended. ganon2 has an advanced custom build procedure with examples for commonly used data sources in the documentation: https://pirovc.github.io/ganon/custom_databases/

16 short-read simulated samples with available ground truth were selected to evaluate the performance of ganon2 and similar tools (Table 2). The 16 samples are very diverse and range from low-to high-complexity environments from different studies, with well-represented organisms (in relation to our chosen reference sets) to poorly represented, with some easier sets (Mende and Microba) to more challenging sets (CAMI1 and CAMI2). For the CAMI sets, we used the first replicate of each study when more than one was available. The goal here is to evaluate all methods using a spectrum of variable diversity and difficulty, instead of performing a one-sample analysis. In this way, methods can be better accessed in different scenarios, and misinterpretation of results can be partially avoided.

**Table 2.**
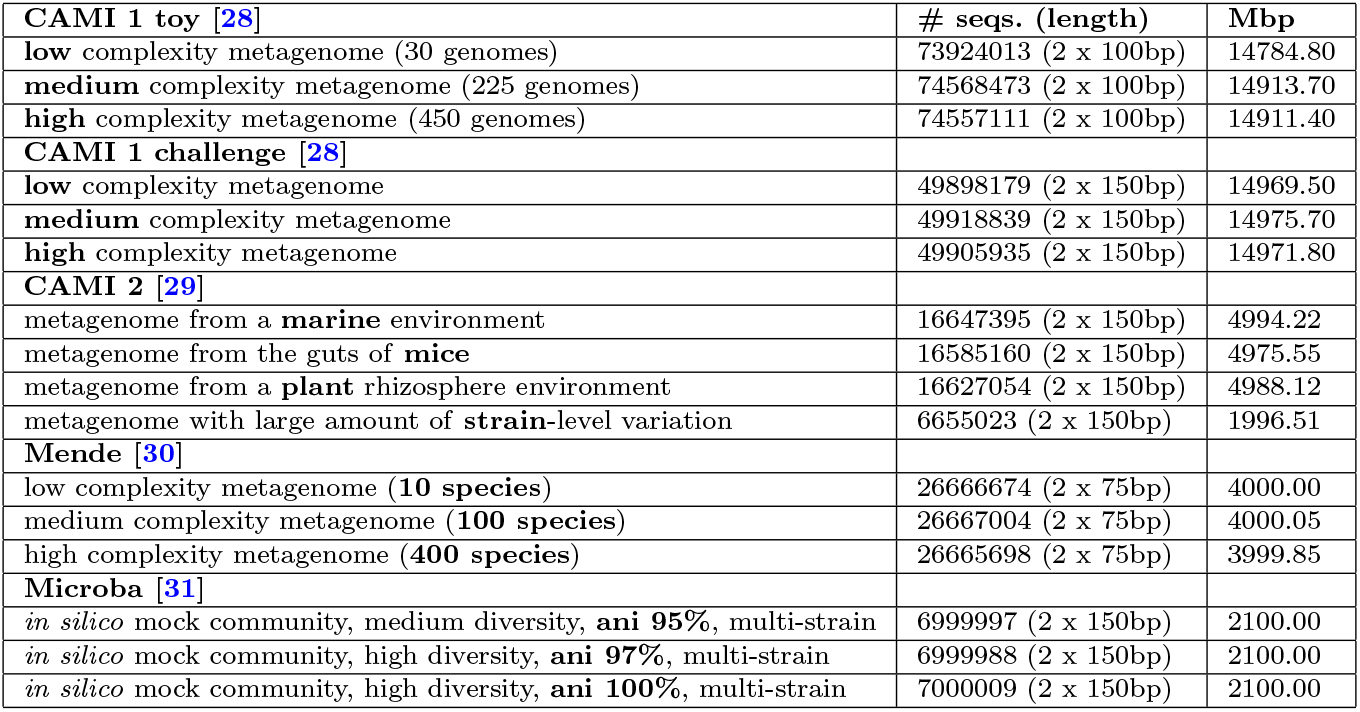
16 simulated short-read samples used in this article, with a total of 124781 Mbp.

|  |  |  |
| --- | --- | --- |
| **CAMI 1 toy [28]** | **# seqs. (length)** | **Mbp** |
| **low** complexity metagenome (30 genomes) | 73924013 (2 x 100bp) | 14784.80 |
| **medium** complexity metagenome (225 genomes) | 74568473 (2 x 100bp) | 14913.70 |
| **high** complexity metagenome (450 genomes) | 74557111 (2 x 100bp) | 14911.40 |
| **CAMI 1 challenge [28]** |  |  |
| **low** complexity metagenome | 49898179 (2 x 150bp) | 14969.50 |
| **medium** complexity metagenome | 49918839 (2 x 150bp) | 14975.70 |
| **high** complexity metagenome | 49905935 (2 x 150bp) | 14971.80 |
| **CAMI 2 [29]** |  |  |
| metagenome from a **marine** environment | 16647395 (2 x 150bp) | 4994.22 |
| metagenome from the guts of **mice** | 16585160 (2 x 150bp) | 4975.55 |
| metagenome from a **plant** rhizosphere environment | 16627054 (2 x 150bp) | 4988.12 |
| metagenome with large amount of **strain**-level variation | 6655023 (2 x 150bp) | 1996.51 |
| **Mende [30]** |  |  |
| low complexity metagenome (**10 species**) | 26666674 (2 x 75bp) | 4000.00 |
| medium complexity metagenome (**100 species**) | 26667004 (2 x 75bp) | 4000.05 |
| high complexity metagenome (**400 species**) | 26665698 (2 x 75bp) | 3999.85 |
| **Microba [31]** |  |  |
| *in silico* mock community, medium diversity, **ani 95%**, multi-strain | 6999997 (2 x 150bp) | 2100.00 |
| *in silico* mock community, high diversity, **ani 97%**, multi-strain | 6999988 (2 x 150bp) | 2100.00 |
| *in silico* mock community, high diversity, **ani 100%**, multi-strain | 7000009 (2 x 150bp) | 2100.00 |

The methods were evaluated in a variety of metrics, with the F1-score being central to summarize the results, since it has the power to evaluate the balance between precision and sensitivity. This balance is very important in metagenomic evaluations, since it is easy to achieve high precision or high sensitivity by classifying just a few or many sequences, respectively. In the binning evaluation, each read is classified to one taxon and set as true or false positive given the ground truth. The number of true and false positives per read classification is then used to calculate the F1-score at a certain taxonomic level (e.g. species). In the profiling evaluation, the F1-Score is based on the final taxonomic profile generated by each method, and the presence of a taxon in the ground truth defines a true positive. The taxa reported that are not present in the truth set are false positives. Additionally, in profiling, we evaluate the abundance estimation for each taxon, and the difference from the ground truth is accounted for in the L1-norm error. For profiling evaluations, a threshold is applied for all results, removing low-abundant taxa. This improves the performance for all methods, reducing the long tail of false positives that are usually reported, and greatly increasing precision with a small decrease in sensitivity. Finally, all methods were evaluated in terms of speed and memory consumption, including custom build of databases. Computational resources used to run all evaluations: 2 x AMD EPYC 9454 48-Core Processor (96 threads), 1TB RAM.

In addition to the self-designed benchmark, we submitted the ganon2 results to the CAMI Portal [32] for the taxonomic profiling and taxonomic binning benchmarks. 3 sets of short-reads challenges [29] were evaluated: Strain-madness (100 samples), plant-associated (21 samples), and marine (10 samples). ganon2 (v2.1.0) was executed against the RefSeq database provided from 2019-01-08, results converted to the required BioBoxes format [33] and submitted to the portal. We then collected the metrics from the website on 2024-03-30. The evaluations here are focused on the metrics provided by the CAMI Portal and the accompanying tools [34–36] and do not include computational performance metrics. They include a broader set of participant methods (e.g. alignment-based, sketching, marker gene-based, protein-based) that do not necessarily use the same reference set or database. Details of the parameters used in each submission can be verified on the CAMI Portal website [32].

During the development of ganon2, we further created a pipeline to run tools, evaluate metrics, and visualize results called MetaBench (https://github.com/pirovc/metabench). This pipeline was used to generate most of the results presented in this article. MetaBench can evaluate taxonomic profiling and binning tools in a combination of databases, parameters, thresholds, and metrics. It outputs standardized file formats (JSON and BioBoxes [33]) and generates a dashboard to visualize them. It was also used to optimize the default parameter configuration for ganon2.

## 3 Results

### 3.1 Build

We first evaluated the performance of the tools to build custom databases, based on 4 RefSeq subsets (Table 1). ganon2 was the fastest method to build the RefSeq RG (20 min), while kmcp was the fastest for the RefSeq CG (14 min) and full RefSeq (6 hours 48 min) (Table 3). kraken2+bracken took the longest to build all databases, between 8 hours and more than 4 days. Metacache has an intermediate time performance, between 1 and a half hours and 12 hours. In terms of memory necessary to build, kmcp has the lowest requirements on average (between 29 and 108 GB) for all databases evaluated, ganon2 coming in second with intermediate memory usage (between 48 and 331 GB). However, kmcp required very large amounts of disk space (several TBs for the full RefSeq) to build the database. bracken and metacache had the highest memory consumption overall, with significantly higher memory usage for the full RefSeq. Finally, when considering database size, ganon2 has, on average, the smallest sizes, with some mixed results for other tools: kmcp has the largest full Ref-Seq database, while kraken2 has the largest for RefSeq RG. Bracken (profiling) is not a standalone tool and depends on the results from kraken2 (binning) to run; therefore, time and database sizes for bracken must be added to kraken2 for realistic numbers. In summary, all tools, besides kraken2 can efficiently build indices for large reference sets in reasonable times, with some bigger differences in memory and disk usage and database size.

**Table 3.** Build time and memory consumption. Time is demonstrated in wall time (lower is better). Memory usage and database size are shown in GB (lower is better). The bracken database is not standalone, and it needs a matching kraken2 database. 64 threads were used for all tools to build the database.

| Time |  |  |  |  |  |
| --- | --- | --- | --- | --- | --- |
|  | ganon2 | kmcp | kraken2 | bracken | metacache |
| RefSeq | 09:47:27 | 06:48:31 | 4 days, 01:29:26 | 02:06:18 | 12:22:11 |
| RefSeq CG | 00:52:44 | 00:14:15 | 11:43:31 | 00:13:51 | 01:37:27 |
| RefSeq RG | 00:20:01 | 00:21:09 | 08:46:14 | 00:09:48 | 01:03:34 |
| RefSeq CG+RG | 01:19:46 | 00:32:22 | 18:50:03 | 00:21:18 | 02:06:37 |
| Memory |  |  |  |  |  |
| (GB) | ganon2 | kmcp | kraken2 | bracken | metacache |
| RefSeq | 331 | 48 | 278 | 505 | 472 |
| RefSeq CG | 78 | 29 | 69 | 83 | 146 |
| RefSeq RG | 146 | 108 | 144 | 157 | 137 |
| RefSeq CG+RG | 168 | 79 | 173 | 188 | 224 |
| Database size |  |  |  |  |  |
| (GB) | ganon2 | kmcp | kraken2 | bracken | metacache |
| RefSeq | 215 | 494 | 274 | 0.019 | 406 |
| RefSeq CG | 42 | 57 | 69 | 0.003 | 119 |
| RefSeq RG | 77 | 34 | 143 | 0.004 | 100 |
| RefSeq CG+RG | 100 | 84 | 172 | 0.006 | 179 |

We also compared the ability and ease of use of tools to generate custom databases. ganon2 provides a highly automated build system. It downloads, builds, and updates any subset of RefSeq and GenBank with only one command, being the most user-friendly tool among the ones tested. Additionally, custom database creation with non-standard files and full integration with NCBI or GTDB taxonomy are also included. kraken2 has the ability to download reference data automatically, but is restricted to fewer options. However, kraken2 does not support compressed input files to build databases, which requires high storage capacities for larger databases, keeping the compressed, uncompressed, and database files on the disk at the same time. Bracken builds upon kraken2 databases. It is straightforward but adds a second step not directly integrated to kraken2 to perform taxonomic profiling. Bracken also requires users to know the size of the reads to be analyzed beforehand, which may require the construction of several databases. kmcp does not automatically download files, but there is an online documentation with guidance on how to perform custom builds. However, the commands are quite extensive and may be a challenge for less experienced users. metacache also has an online documentation on building custom databases, which is well explained but adds a level of complexity that may confuse less technically inclined users.

### 3.2 Binning

The binning results in terms of F1-Score at the species level are presented in Figure 1. ganon2 shows a significant improvement over other methods, especially when using more diverse databases (full RefSeq, RefSeq CG, and RefSeq CG+RG). For the full RefSeq, ganon2 achieved a median F1-score of 0.77, with 75% of the samples with an F1-score of 0.53 or higher, with a well-balanced performance between precision and sensitivity, as shown in Figure 2. kraken2 comes second with a median of 0.61, followed by metacache with 0.45. kmcp could not be evaluated with the full RefSeq due to very long run times. The same trend with slightly lower values can be observed for RefSeq CG+RG, followed by RefSeq CG, both providing very good results with significantly smaller databases (Table 1). With the smallest set of references (RefSeq RG), tools have similar results, with a median of around 0.55. Metacache performed badly in this sample, having most of the assignments to lower ranks (e.g. genus, family). Those results show how tools can perform drastically differently with mixed databases and the benefits of using a highly diverse and larger set of references. ganon2 has the smallest memory footprint for classifying reads against all databases in addition to kmcp which just finished the run for the smallest reference set (Table 4). Using RefSeq CG, ganon2 also achieved very good results in terms of F1-score and requires 5.1 times less memory (42 GB) compared to the full RefSeq (217 GB). metacache was the fastest tool to perform binning; however, it had higher memory usage and poor results in this scenario. kraken2 and ganon2 have similar performance in terms of speed and kraken2 is significantly faster with the full RefSeq. ganon2 can also take advantage of multiple threads to improve run-times (Supplementary Figure 1). kmcp has the smallest memory footprint using the RefSeq RG, but was the slowest among the tested methods, failing to provide results in a doable time (several days) for any of the largest databases.

**Table 4.** Binning and profiling speed and memory consumption, averaged from 16 short-read samples for every tool and database combination. Speed is demonstrated in Mbp/m (higher is better) with standard deviation in brackets. Peak memory is shown in GB. Every combination of tool, database, and sample was executed 3 consecutive times, and only the fastest was considered. The values represent a full run, including pre- and post-processing steps (loading the database in memory, classification, read reassignment, etc.). kmcp did not finish the benchmark after several days with the full RefSeq, RefSeq CG, and RefSeq CG+RG databases; therefore it is not included. 64 threads were used.

|  |  |  |  |  |
| --- | --- | --- | --- | --- |
| **Binning** |  |  |  |  |
| Speed (Mbp/m) | **ganon2** | **kmcp** | **kraken2** | **metacache** |
| **RefSeq** | 1114 ($\pm 527$) | - | 2237 ($\pm 591$) | 1296 ($\pm 683$) |
| **RefSeq CG** | 2523 ($\pm 765$) | - | 2908 ($\pm 611$) | 4640 ($\pm 1192$) |
| **RefSeq RG** | 2345 ($\pm 547$) | 483 ($\pm 85$) | 2127 ($\pm 555$) | 5041 ($\pm 930$) |
| **RefSeq CG+RG** | 2196 ($\pm 657$) | - | 2285 ($\pm 688$) | 3603 ($\pm 1070$) |
| Mem. (GB) | **ganon2** | **kmcp** | **kraken2** | **metacache** |
| **RefSeq** | 217 | - | 279 | 435 |
| **RefSeq CG** | 42 | - | 72 | 124 |
| **RefSeq RG** | 78 | 34 | 145 | 108 |
| **RefSeq CG+RG** | 100 | - | 177 | 187 |
| **Profiling** |  |  |  |  |
| Speed (Mbp/m) | **ganon2** | **kmcp** | **kraken2+bracken** |  |
| **RefSeq** | 1664 ($\pm 937$) | - | 2218 ($\pm 591$) | |
| **RefSeq CG** | 3007 ($\pm 726$) | - | 2892 ($\pm 605$) | |
| **RefSeq RG** | 2868 ($\pm 746$) | 434 ($\pm 103$) | 2118 ($\pm 556$) | |
| **RefSeq CG+RG** | 2734 ($\pm 755$) | - | 2272 ($\pm 685$) | |
| Mem. (GB) | **ganon2** | **kmcp** | **kraken2+bracken** |  |
| **RefSeq** | 216 | - | 279 |  |
| **RefSeq CG** | 42 | - | 72 |  |
| **RefSeq RG** | 78 | 34 | 145 |  |
| **RefSeq CG+RG** | 100 | - | 177 |  |

**Fig. 1.**
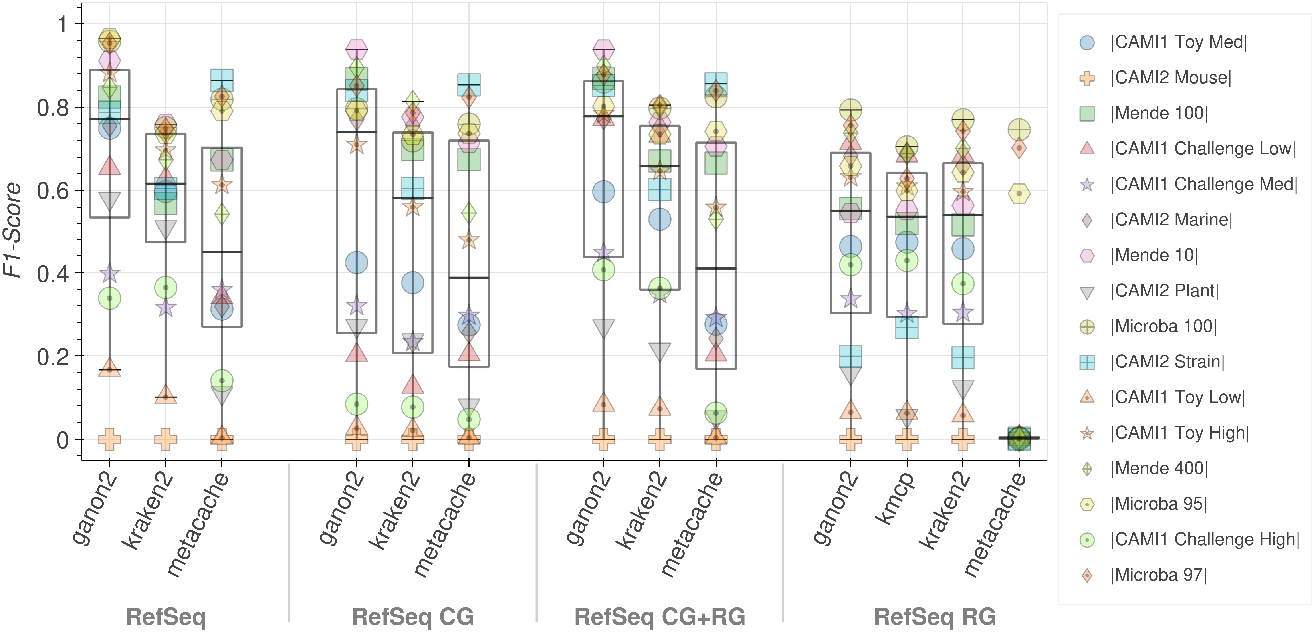
F1-Score results for taxonomic binning at the species level (higher is better). Each tool and database combination (x-axis) has 16 points, one for each sample analyzed (Table 2), identified by a distinct marker and color, with a boxplot showing their overall distribution. The CAMI 2 mice dataset (cross) scored 0 for all methods because the ground truth is provided at the genus level.

**Fig. 2.**
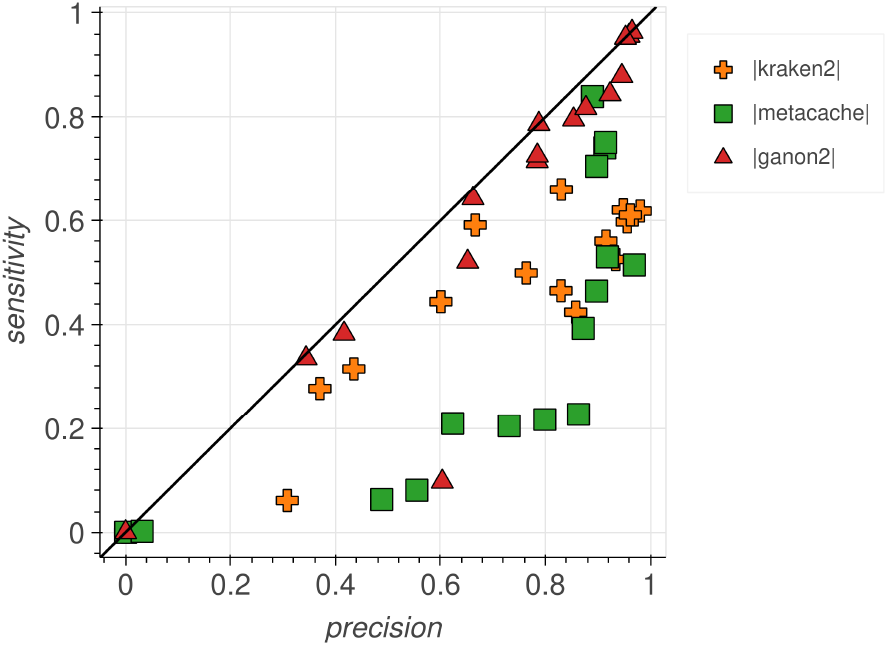
Sensitivity and precision plot for taxonomic binning at the species level against the full RefSeq. Each tool has 16 points, one for each sample analyzed (Table 2). ganon2 achieved balanced results between the metrics.

### 3.3 Profiling

In profiling, ganon2 shows a highly improved F1-score at the species level with the full RefSeq, with 50% of the samples with an F1-score above 0.67 (Figure 3 TOP). In the same scenario, kraken2 + bracken achieved a median of 0.31. kmcp shows comprable good results to ganon2 with the RefSeq RG databases; however, it takes on average at least 6 times longer to run (Table 4). To illustrate the difference in runtime using the RefSeq RG in profiling: ganon2 took 43 minutes to run all 16 samples (Table 2) in this benchmark, kraken2+bracken took 59 minutes, while kmcp needed 4 hours and 47 minutes. Although fast, kraken2 + bracken performed poorly in our profiling evaluation in terms of F1-score. Here again, the results clearly show the benefits of using a larger database (full RefSeq and RefSeq CG+RG) to profile diverse samples, which results in a substantial improvement. With diverse databases, ganon2 runs fast, with the smallest memory footprint, and achieves the best performance in terms of F1-Score among 16 samples.

**Fig. 3.**
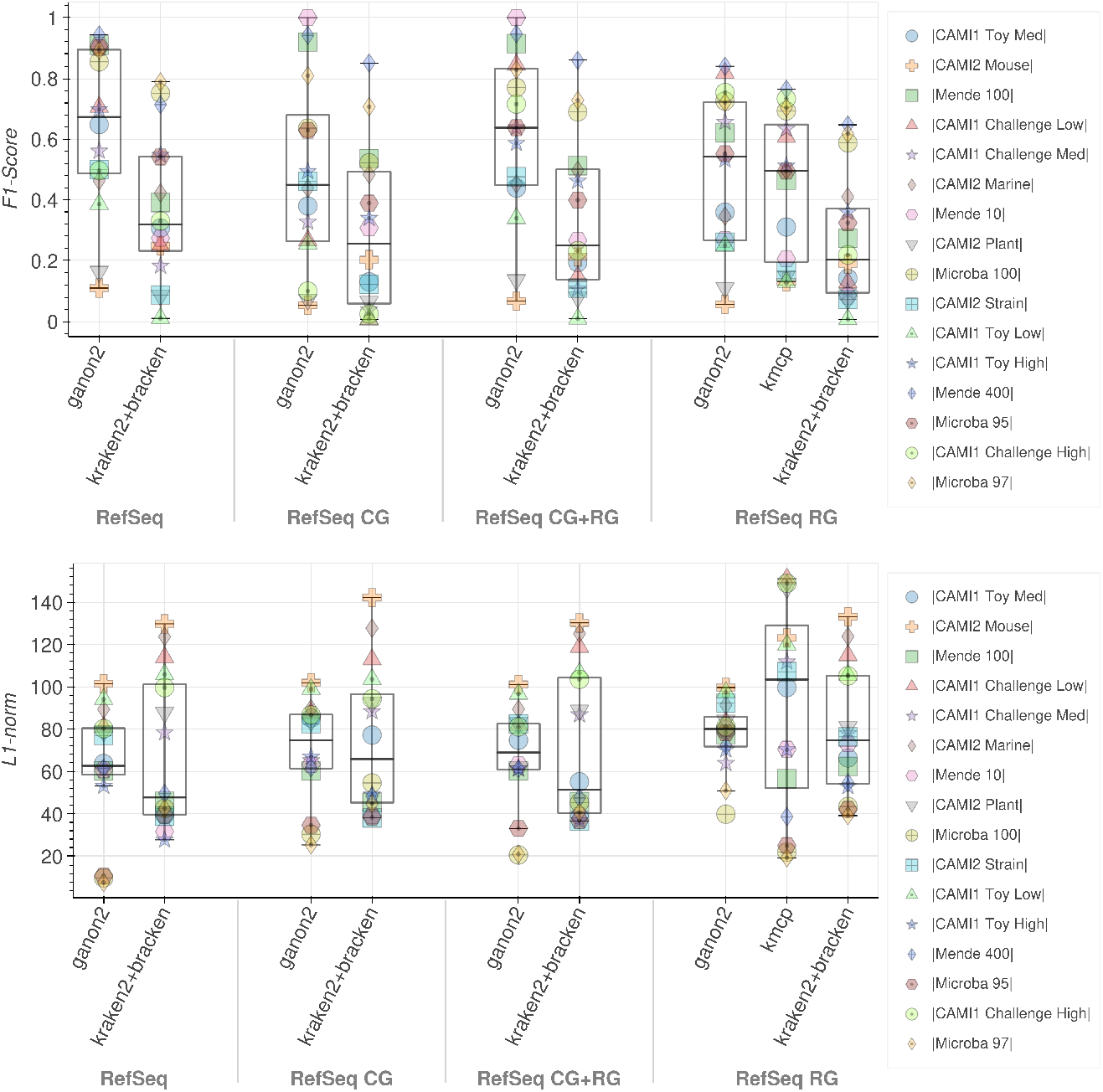
**TOP**) F1-Score results for taxonomic profiling at the species level (higher is better). **BOT-TOM**) L1-norm error results at the species level (lower is better). Each tool and database combination (x-axis) has 16 points, one for each sample analyzed (Table 2), identified by a distinct marker and color, with a boxplot showing their overall distribution. Results considering only species with abundance above 0.005% for all tools.

The L1-norm error analysis shows how the tools perform in terms of abundance estimation (Figure 3 BOTTOM). Here, tools behave quite differently, with ganon2 showing the smallest interquartile range in all scenarios, followed by kraken2 + bracken and kmcp. kmcp always reports a full 100% abundance in the taxonomic profiles, not considering unmatched reads or unknown taxa. This is beneficial when samples are well represented and covered primarily by the database (Microba and Mende sets) but can generate inflated abundances when that is not the case (CAMI sets), as shown in Figure 3. kraken2+bracken also has a higher number of classified reads, which explains the higher variation in L1-norm values. In all scenarios, ganon2 did not exceed L1-norm values of approximately 100 for all samples, keeping a low error in abundance estimations. kmcp has very low L1-norm values for some samples with RefSeq CG, with the downside of also having some of the highest L1-norm for others.

Further metrics, thresholds, and plots for the results presented above can be interactively explored in the Supplementary Material HTML report.

### 3.4 CAMI Portal bechmarks

The results presented in this section were directly extracted from the CAMI Portal on 2025-03-30. Figures 4 - 5 show the trade-off between purity (precision) and completeness (sensitivity) at the species level, as well as the list of tools ranked from best to worst for each data set. The results are based on the average of samples for each set (100 strain-madness samples, 21 plant-associate samples, 10 marine samples) for several methods: Bracken [25], CCMetagen [37], Centrifuge [38], Centrifuger [39], DIAMOND [40], DUDes [41], ganon [13], kraken2 [24], LSHVec [42], Metabuli [43], MetaPhlAn [44], mOTUs [45], NBC++ [46], sourmash [47], Sylph [48], TIPP [49]. ganon2 scores well in all CAMI2 short-read challenge sets for taxonomic profiling

**Fig. 4.**
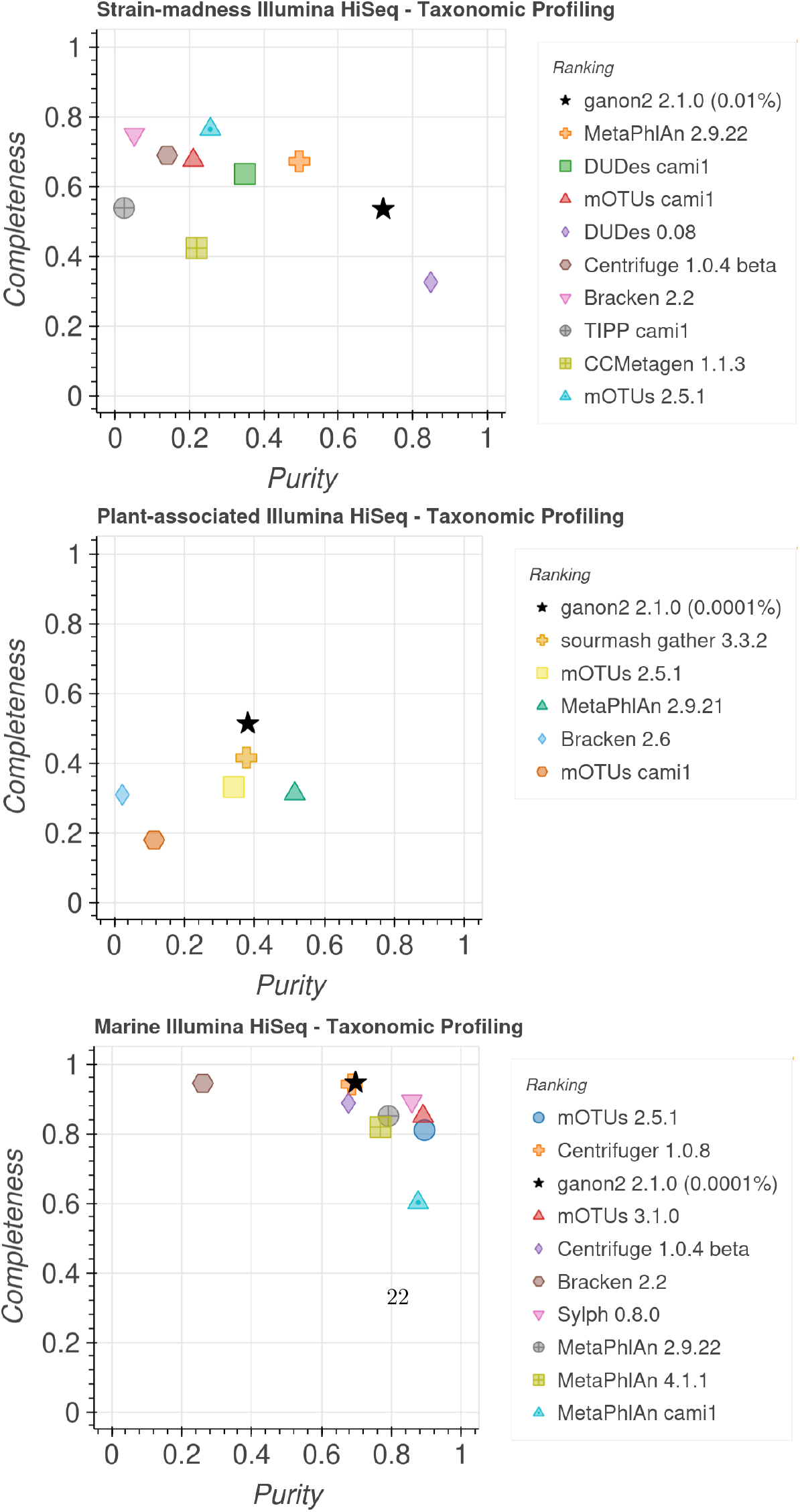
Completeness (sensitivity) and Purity (precision) plot for taxonomic profiling at the species level. Data extracted from the CAMI portal [32] 2025-03-30, based on the average over all samples for each set. Only the best result for each combination of tool + versions was kept and the top 10 results for each set are presented. Tools are sorted in the legend based on their ranking which is generated based on additional metrics not displayed. Values in parenthesis in the legend show the minimum abundance threshold used for ganon2 results.

(Figure 4). It ranked first in the strain-madness and plant-associated sets and third in the marine set. The ranking incorporates additional data not displayed (completeness, purity, F1 score, L1 norm error, Bray-Curtis distance, Shannon equitability, and weighted UniFrac error) and shows that ganon2 can achieve overall good and balanced results for various sample characteristics. Marker gene-based tools (mOTUs and MetaPhlAn) achieved good purity and completeness results, but in some sets were ranked low because of lower values on other metrics. sourmash and Centrifuger achieved good results in specific sets (plant-associated and marine, respectively); however, data is not available for the other sets, so they can only be partially evaluated. Similarly, for the taxonomic binning challenge (Figure 5) ganon2 ranked first in the Marine set, in line with our own evaluations. Other sets for the taxonomic binning challenge have fewer or no other methods submitted, and therefore, were omitted here. More results with all metrics, ranks, and CAMI1 challenge sets are available at https://cami-challenge.org/ and in the Supplementary Material HTML reports.

**Fig. 5.**
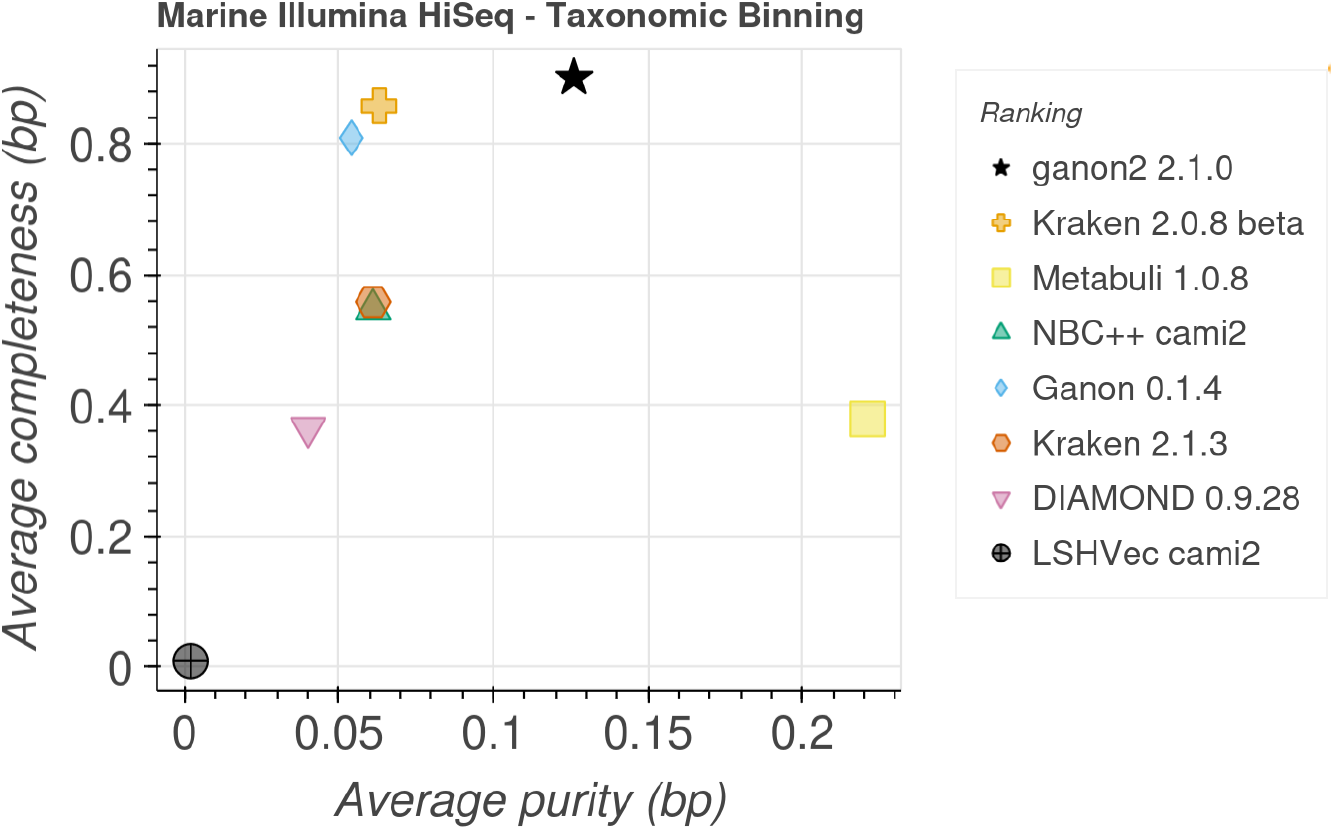
Completeness (sensitivity) and Purity (precision) plot for taxonomic binning at the species level. Data extracted from the CAMI portal [32] on 2025-03-30. Only the best result for each combination of tool + versions was kept. Tools are sorted in the legend based on their ranking which is generated based on additional metrics not displayed.

### 3.5 Sewage metagenomics analysis

To demonstrate ganon2 use in a real-world scenario, we re-analyzed sewage metagenomics samples. The original study [50] analyzed time-series of sewage samples from several European cities, with the aim of understanding the data patterns and developing a quantification and correlation workflow to monitor antimicrobrial resistance over time.

ganon2 was used to classify the trimmed raw reads of the 28 samples from the city of Bologna, Italy (data available at (https://www.ebi.ac.uk/ena/browser/view/PRJEB68319). As a reference, the entire GTDB R220 (596859 genomes) was used [51]. We then compared the short-read ganon2 output with the taxonomic profile provided by the study ([50] Figure 1b). ganon2 achieved comparable abundances at the genus level by directly analyzing short-reads, without the need of genome assembly, as demonstrated in Figure 6. Blooms of *Pseudomonas E* were correctly detected and most of the reported genus were closely profiled. This not only reproduces and confirms the findings and methods of the original study, but also highlights the ability of ganon2 to properly and quickly profile microbiome samples accurately. This can be crucial in real-time genomic surveillance [52] (e.g. pathogen detection [53]), since genome assembly is a computationally heavy and time-consuming process, as stated by the authors of the sewage study, especially in co-assembly for metagenomics: “A downsize was the extensive compute requirements with some jobs running for many weeks and needing a lot of memory.” [50]. Metagenomic assembly is an essential technique for in-depth and reference-free analysis of microbiome samples and is crucial to improve public collections of references from unknown environments. However, short-read metagenomics screenings provide a quick alternative to the initial analysis and profile of environmental samples with a fraction of the time and computational cost.

**Fig. 6.**
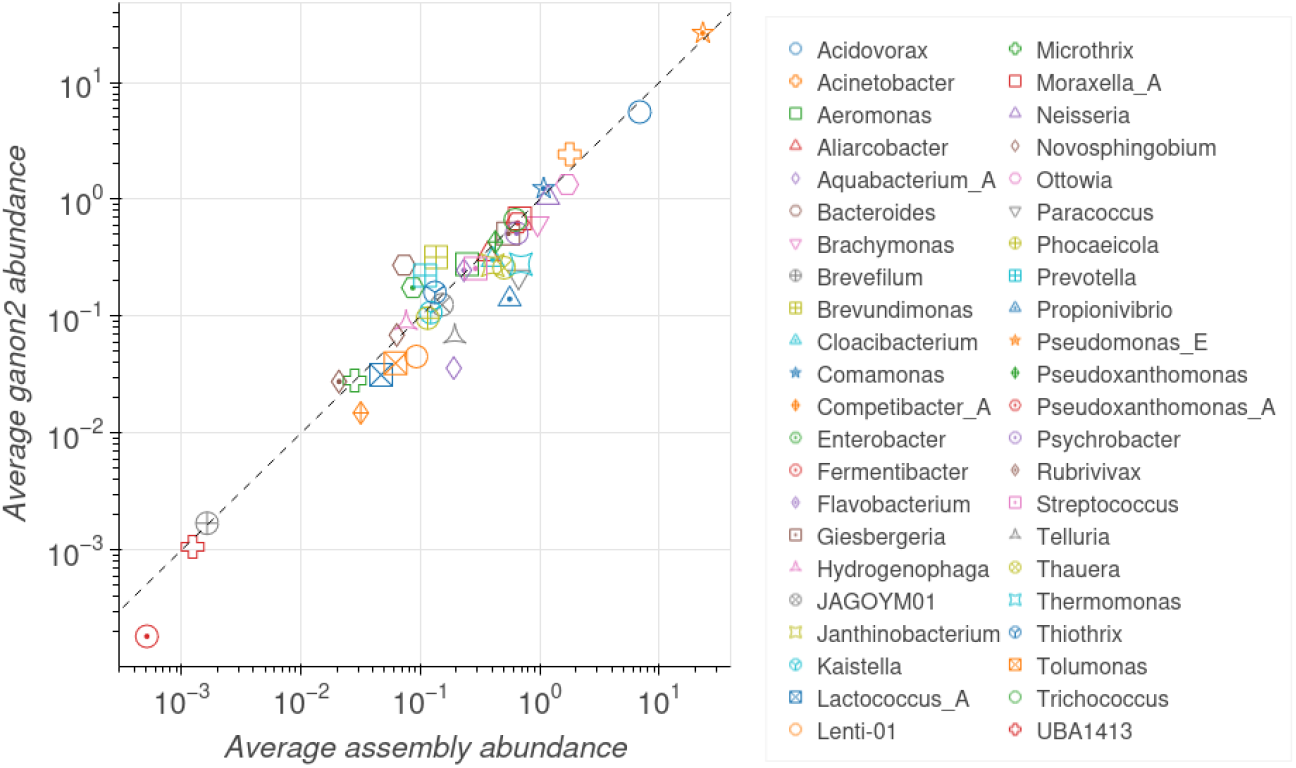
Average abundances among the 28 samples from Bologna, Italy for each of the 44 genus reported by [50] based on assembled data compared to the normalized averages obtained with ganon2 based on short-read data. Axis are in log-scale. The dotted line is the diagonal slope used for reference.

## 4 Discussion and Conclusion

ganon2 performs metagenomics analysis, integrating database download and construction with binning and profiling classification methods. It further brings an extensive set of exclusive functions: multiple and hierarchical database support, customizable database construction for local or non-standard sequence files, NCBI taxonomy and GTDB integration, optional LCA or EM-algorithm to solve multi-matching reads, classification of short and long reads, optimized taxonomic binning and classification configurations, advanced report generation and filtering, and easy integrated analysis at any taxonomic rank, strain, assembly, file, sequence, or custom specialization level. Coupled with a set of simple parameters and extensive documentation, ganon2 is a robust tool for exploratory metagenomic classification and data analysis.

Using 16 diverse, realistic, and complementary samples against 4 different genomic reference sets based on RefSeq, ganon2 achieved consistently better results than comparable methods while maintaining top performance in terms of runtime and memory consumption. Although the tools evaluated perform similarly well using standard size databases, ganon2 excels when using more data, scaling well. This gives more options for researchers, especially when describing under-represented environments, where a combination of many resources is necessary to minimally profile samples. ganon2 is able to classify more reads correctly, increasing the resolution of the results. In addition to our self-designed evaluation, results were uploaded to the CAMI Portal, where they were independently evaluated based on a set of various metrics against many tools. ganon2 achieved very competitive results, ranking first in many of the challenge sets.

By default, a minimum abundance threshold was applied for all methods in our benchmarks. This greatly improves precision metrics, since all methods generate a long tail of false positives results based on taxa that received only a few matches. The decision on this threshold value can be arbitrary, since each sample is different and low-abundance organisms may mix with false signals. As a rule of thumb and based on observations during our benchmarks, the use of any, even a very low threshold, is already advantageous and should be always applied. For the CAMI2 Portal results, we applied different thresholds based on the known number of organisms to be detected in each sample.

The results in this article are focused on species level. However, ganon2 can build indices at any taxonomic level or at more specific targets such as assembly, strain, file, and sequence. Different levels are treated equally by the ganon2 algorithm and strain classification can be performed, but cutoff and filter threshold values should be adjusted to obtain more sensitive or precise results. Specific parameterization and optimization for strain-level classification could be a fruitful future research.

The improvements in ganon2 were achieved with a combination of factors. First, the use of the Hierarchical Interleaved Bloom Filter paired with minimizers over the Interleaved Bloom Filter improved performance and at the same time reduced drastically database sizes and memory requirements. Second, an extensive optimization of parameters and methodologies based on real data for the post-processing procedure (e.g. read reassignment) refined the results, enabling ganon2 to achieve a strong overall improvement in evaluation metrics compared to its first version and the current methodologies.

We are aware of and acknowledge the difficulties and biases in evaluating self-developed methods in an article fairly versus others [54]. In our evaluations, we ran all methods in default mode using the same underlying set of reference sequences. We believe that this is the fairest way to evaluate methods in a user-level approach. As noted in other studies [9, 29], changing parameters on a sample basis can drastically change the results of methods. However, real users in a real-case application usually do not have the time and resources to fine-tune the parameters for every analysis and do not have the ground truth to evaluate the effects of the changes. Observing the usage of such methods in the literature, in most cases they are executed with the provided default parameters with little to no variation. Therefore, in our opinion, this is the most sensible way to evaluate them.

On a more technical level, many methods share characteristics in terms of data structure and parameterization (e.g. k-mer size), but the equivalence of them can be complex and not directly comparable. For example, kmcp uses a variation of bloom filters with a false-positive rate of 30% and one hash function by default, while ganon2 also uses variations of bloom filters, but with a lower false-positive rate and more hash functions. The way both tools use the outcome from the bloom filters is different, and matching those values would not make the comparison fairer. The reader should be aware of possible bias in software articles and interpret the results with discretion [54]. We mainly benchmarked tools that employ similar methods, based on k-mer complete genomic comparisons, therefore sharing most of the computational trade-offs. There are currently a diverse set of tools to perform similar tasks based on different techniques, e.g.: marker-gene, protein, alignment, and sketching-based, among others. Each of them has their own trade-offs in terms of computational resources and classification performance. Furthermore, the choice of database and tool parameterization plays a crucial role in the computational performance and results. A fully fledged comparison between all of these variables can be extremely complex and is out of the scope of this article. Other benchmark articles [55] and the CAMI Portal benchmark provide an idea on how different methods compare, at least in terms of classification results.

In an effort towards transparency and reproducibility, we provide all results and metrics from this article in an easy-to-navigate, open-source, and user-friendly dashboard (Supplementary Material HTML report), where it is possible to verify the results of each metric for all samples, databases, and methods independently and how they were performed. A completely fair and unbiased evaluation can only be performed by researchers who are not related to the development of the evaluated methods. Independent and external evaluation efforts such as LEMMI [56] and CAMI [29] are great resources; however, a continuous, updated, and actively supported benchmark using the capabilities of MetaBench, covering a range of parameter variations and databases, is still to be developed.

ganon2 provides an efficient and scalable method to analyze large amounts of sequences against numerous reference sequences. It excels in metagenomics analysis, providing substantial improvements in taxonomic binning and profiling, especially using larger and more diverse reference sets. ganon2 is fast in terms of building databases and analyzing data, with a reasonable memory footprint and the best results in terms of sensitivity and precision in our simulated evaluations. Not only does it provide an improvement over its first version, but it is also the overall best-performing tool in comparison with some of the state-of-the-art methods. ganon2 was developed with ease of use in mind, allowing the use of diverse, large, and up-to-date reference sequences in metagenomic analysis. ganon2 is open-source and comes with extensive documentation online with a detailed explanation of its functions, databases, parameters, and trade-offs and can be found at https://pirovc.github.io/ganon/.

## Supporting information

Supplementary Material HTML report

## Data Availability

The source code for ganon2 can be found in GitHub (https://github.com/pirovc/ganon) and Zenodo (v2.1.0: https://doi.org/10.5281/zenodo.10648995). The source code for MetaBench can be found in GitHub (https://github.com/pirovc/metabench) and Zenodo (v1.1.0: https://doi.org/10.5281/zenodo.14330683).

