## Supplementary Figures for "ganon2: up-to-date and scalable metagenomics analysis"

### ganon2: up-to-date and scalable metagenomics analysis
### Supplementary Figures

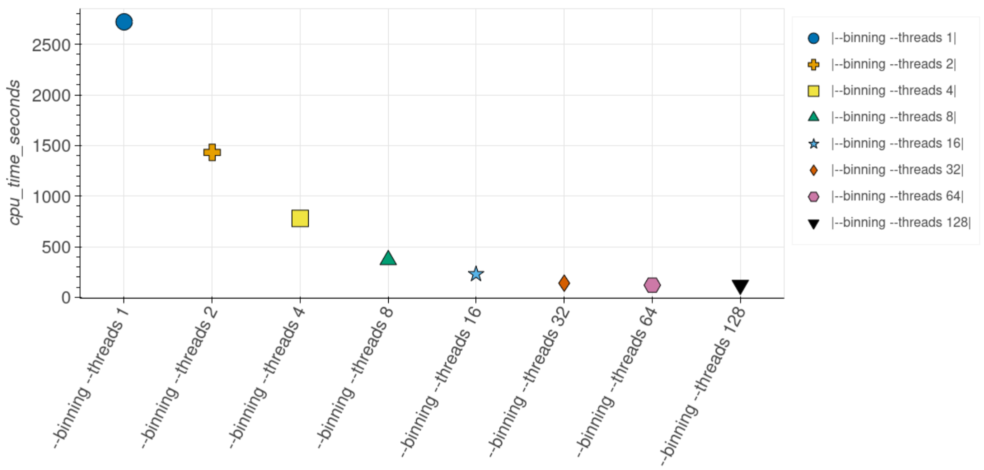


**Supplementary Figure 1** Time (seconds) to profile one CAMI2 Challenge Marine sample against the RefSeq CG+RG with different number of threads. Each parametrization was executed 3 consecutive times and only the fastest was considered.
